# Changes in free amino acid concentrations and associated gene expression profiles in the abdominal muscle of kuruma shrimp *Marsupenaeus japonicus* reared at different salinity

**DOI:** 10.1101/180638

**Authors:** Hiroki Koyama, Nanami Mizusawa, Masataka Hoashi, Enkong Tan, Ko Yasumoto, Mitsuru Jimbo, Daisuke Ikeda, Takehiko Yokoyama, Shuichi Asakawa, Sanit Piyapattanakorn, Shugo Watabe

**Affiliations:** Graduate School of Biosphere Science, Hiroshima University, Hiroshima 739-8528, Japan; Kitasato University School of Marine Biosciences, Kanagawa 252-0373, Japan; Graduate School of Agricultural and Life Sciences, The University of Tokyo, Tokyo 113-8657, Japan; Department of Marine Science, Faculty of Science, Chulalongkorn University, Bangkok 10330, Thailand

**Keywords:** Free amino acids, Kuruma shrimp, Marsupenaeus japonicus, RNA-seq, Salinity

## Abstract

Shrimps inhabiting around the coastal area can survive in a wide range of salinity. However, the molecular mechanisms involved in their adaptation to different environmental salinity have remained largely unknown. In the present study, we reared kuruma shrimp *Marsupenaeus japonicus* at 1.7 %, 3.4 % and 4.0 % salinity. After rearing for 6, 12, 24 and 72 h, we determined free amino acid concentrations in their abdominal muscle, and performed RNA-seq analysis on this muscle. The concentrations of free amino acids were clearly altered depending on salinity after rearing for 24 h. Glutamine and alanine concentrations were markedly increased following the increase of salinity. In association with such changes, many genes related to amino acid metabolism changed their expression levels. Notably, the increased glutamine content at high salinity appeared to be relevant to the increase of the expression level of the gene encoding glutamate-ammonia ligase which functions in the glutamine metabolism. Furthermore, the alanine content increased at high salinity was likely to be associated with the decrease in the expression levels of the alanine-glyoxylate transaminase gene. Thus, the changes in the concentration of free amino acids for osmoregulation in kuruma shrimp are considered to be regulated by the changes in the expression levels of genes related to amino acid metabolism.

**Summary statesment:** Kuruma shrimp *Marsupenaeus japonicus* changes free amino acid contents and associated gene expression levels in their muscle to adjust effectively to different salinity.

## Introduction

Shrimps belong to the class Crustacea, which form a large diverse group in invertebrates and some of them have exploited their niche by adaptation to different temperature as an isolation factor (David, 2014; Jorde *et al.*, 2015; Martin and Davis, 2001). Several shrimps inhabiting the coastal area can survive in a wide range of salinity, by changing intracellular free amino acid concentrations to maintain osmotic pressures (Camien *et at.*, 1951; Freire *et al.*, 2008; Henry *et al.*, 1980; McNamara *et al.*, 2004). It has been reported that the concentrations of total free amino acids were increased in the muscles of crayfish *Procambarus clarkii* and kuruma shrimp *Marsupenaeus japonicus* following the increase of environmental salinity, and the changes were largely due to those of glycine and L-alanine (Abe *et al.*, 2005; Okuma and Abe, 1994). Therefore, the two amino acids are considered to be important osmolytes for these invertebrates (Abe *et al.*, 1999, 2005; Fujimori and Abe, 2002; Okuma and Abe, 1994). Another experiments indicated that the concentrations of total free amino acids were decreased in the muscle of Pacific white shrimp *Litopenaeus vannamei* reared at low salinity, whereas those of glycine and L-serine were increased in the hemolymph to decrease the osmotic pressure in the muscle, suggesting that tissue amino acids were released into the hemolymph to lower the osmolarity of the tissue (Shinji *et al.*, 2012).

Despite such fact, the adaptation to different salinity will be alternatively regulated by the particular genes. Suppression subtractive hybridization and real-time PCR revealed the relationship between environmental salinity and gene expression levels. For instance, black tiger shrimp *Penaeus monodon* (Shekhar *et al.*, 2013, 2014), Pacific white shrimp (Gao *et al.*, 2012; Sun *et al.*, 2011) and ridgetail white prawn *Exopalaemon carinicauda* (Li *et al.*, 2015) increased the expression levels of the gene encoding Na^+^/K^+^-ATPase α-subunit in various tissues such as gills, gut, hepatopancreas and antennal glands exposed to either high or low salinity. Na^+^/K^+^-ATPase is known as one of the ion transporters which exchange ions between cytoplasm and hemolymph to maintain inorganic ion concentrations in shrimp (Boudour-Boucheker *et al.*, 2014; Faleiros *et al.*, 2010; Havird *et al.*, 2014; Holliday, 1985), suggesting that Na^+^/K^+^-ATPase plays an important role in osmoregulatory systems at both high and low salinity. Black tiger shrimp exposed to high salinity also increased the expression levels of genes encoding intracellular fatty acid binding proteins in gut tissues (Shekhar *et al.*, 2013), whereas Pacific white shrimp decreased those encoding hemocyanin, chitinase, ecdysteroid-regulated protein, trypsin and chymotrypsin 1 in hepatopancreas (Gao *et al.*, 2012; Sun *et al.*, 2011).

RNA-seq analysis has been demonstrated to be a powerful method to examine the effects of salinity or temperature on gene expression levels in several invertebrates (Huang *et al.*, 2017; Lv *et al.*, 2013; Meng *et al.*, 2013; Santos *et al.*, 2014; Sellars *et al.*, 2015; Zhao *et al.*, 2012). It has been reported that swimming crab *Portunus trituberculatus* reared for ten days at different salinity changed the expression levels of osmoregulation-related genes such as those encoding ion transporters and amino acid metabolism-related proteins in their gills (Lv *et al.*, 2013).

As mentioned above, many genes including ion and amino acid transporters seem to participate in adaptation of crustaceans to the salinity change. However, the molecular mechanisms involved have still remained unclear, since the regulatory mechanisms underlying the changes of free amino acid concentrations are not well understood.

In the present study, we targeted kuruma shrimp as experimental animals, which are widely cultured and commercially available, important species. We reared the shrimp samples at different salinity and determined free amino acid concentrations in their abdominal muscle. In addition, we performed RNA-seq analysis on the same samples by using a next generation sequencer.

## Materials and methods

### Animals

About 40 adult specimens of kuruma shrimp were obtained from Miyazaki Prefecture, Japan, and cultured in 3.4 % salinity tank (60 l) of Kitasato University at 25 °C for 3 days. Then, they were divided into 3 groups using 60 l tanks each at 1.7 %, 3.4 % and 4.0 % salinity. Shrimp were fed commercially available pellets for shrimp ad libitum under about 14 h: 10 h light: dark cycle. After rearing at 25 °C for 6, 12, 24 and 72 h, three specimens were collected each from the three tanks. The body lengths and weights of kuruma shrimp samples are shown in Table 1.

**Table 1.** Body length and body weight of kuruma shrimp reared at different salinity

| Salinity | 0 h |  | 6 h |  | 12 h |  | 24 h |  | 72 h |  |
| --- | --- | --- | --- | --- | --- | --- | --- | --- | --- | --- |
|  | Body length (cm) | Body weight (g) | Body length (cm) | Body weight (g) | Body length (cm) | Body weight (g) | Body length (cm) | Body weight (g) | Body length (cm) | Body weight (g) |
| 1.7% |  |  | 12.2 ± 0.21 | 15.5 ± 0.28 | 11.1 ± 0.38 | 12.1 ± 1.45 | 11.8 ± 0.16 | 10.9 ± 0.33 | 12.7 ± 0.21 | 15.3 ± 0.82 |
| 3.4% | 10.9 ± 0.37 | 9.9 ± 0.91 | 12.4 ± 0.24 | 15.0 ± 0.56 | 11.2 ± 0.21 | 11.1 ± 0.41 | 11.4 ± 0.37 | 11.5 ± 0.34 | 12.4 ± 0.08 | 15.5 ± 0.61 |
| 4.0% |  |  | 12.4 ± 0.05 | 15.3 ± 0.33 | 11.9 ± 0.05 | 11.9 ± 0.33 | 11.7 ± 0.19 | 13.6 ± 0.37 | 12.2 ± 0.24 | 15.8 ± 0.54 |

Values are given as mean ± standard deviation (n = 3).

### The determination of free amino acid concentrations

The second abdominal segments (Fig. 1) of three kuruma shrimp each from different salinity tanks were dissected after rearing for different periods, homogenized individually with 8 volumes of 10 % perchloric acid (w/w), and centrifuged at 12,000 *g* for 20 min at 4 °C. The resulting supernatant was neutralized with an appropriate amount of 12 mol l^-1^ and 2 mol l^-1^ KOH, and centrifuged at 12,000 *g* for 20 min at 4 °C to collect the supernatant containing free amino acids. Free amino acids were derivatized with *O*-phthalaldehyde and 3-mercaptopropionic acid, and their concentrations were determined by using a high performance liquid chromatography LC-2000 series (Jasco, Tokyo, Japan) with a reverse-phase column TSK gel ODS-80Ts (length, 250 mm; inside diameter, 46 mm) (Tosoh, Tokyo, Japan). Mobile phase A consisted of 50 mmol l^-1^ sodium acetate buffer (pH 5.63) and mobile phase B, 20 % 50 mmol l^-1^ sodium acetate buffer (pH 5.63) plus 80 % absolute methanol (v/v). Amino acids were eluted at room temperature with a linear gradient from A: B = 100: 0 to A: B =10: 90 in 75 min at a flow rate of 1.0 ml min^-1^. Excitation and emission wavelengths to detect derivatized amino acids were 340 nm and 450 nm, respectively. The present method cannot distinguish between L- and D-amino acids.

**Fig. 1.**
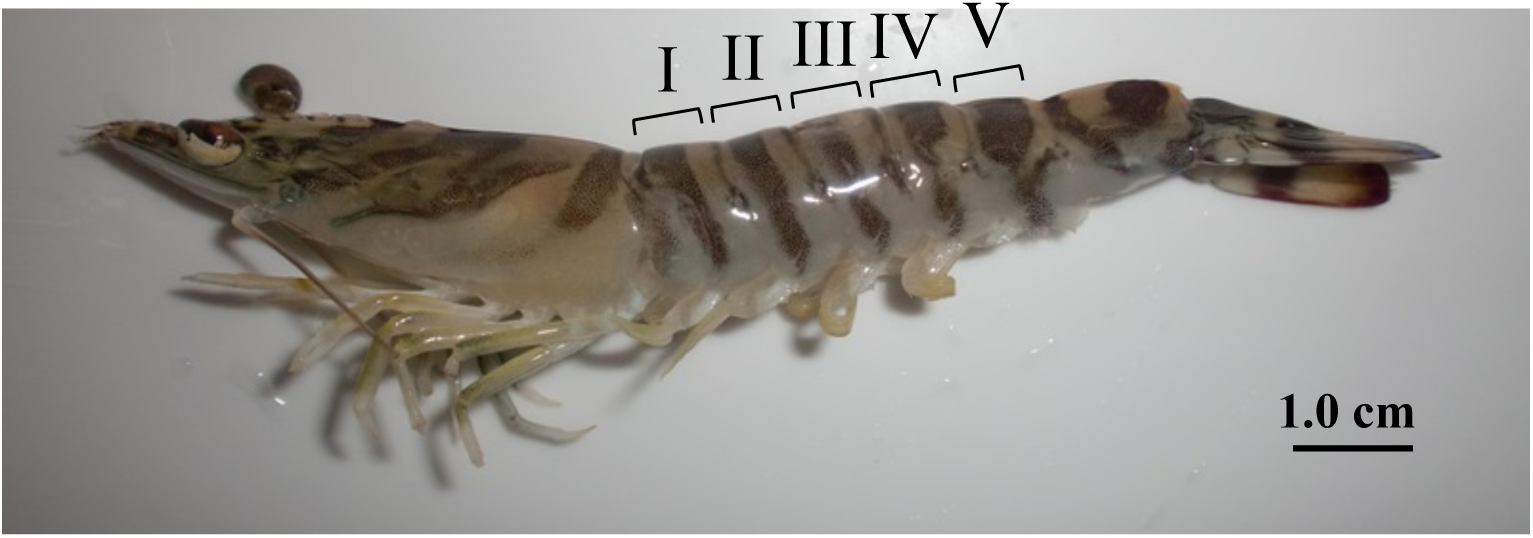
The figure of kuruma shrimp. I-V correspond to the number of individual abdominal segments.

### Water content

The fourth and fifth abdominal segments (Fig. 1) of three specimens each from different salinity tanks were collected after rearing for various periods and minced individually. The water content was measured with an MA35 moisture meter (Sartorius, GLttingen, Germany) according to the manufacturer’s instructions.

### The construction of cDNA library

Total RNAs were extracted from the third abdominal segments (Fig. 1) each of three specimens reared at different salinity for 24 h, using ISOGEN II solution (Nippon Gene, Tokyo, Japan) according to the manufacturer’s instructions, where 30 μg of total RNAs each from three specimens reared at 1.7 %, 3.4 % or 4.0 % salinity tanks were mixed, respectively. These total RNAs were treated with DNase I (Takara, Otsu, Japan) to digest contaminated gDNA. Then, mRNAs were purified by using Poly(A)^+^ Isolation Kit from Total RNA (Nippon Gene). Subsequently, cDNA libraries were constructed from purified mRNAs by using Ion Total RNA-Seq Kit v2 (Life Technologies, Carlsbad, CA, USA) according to the manufacturer’s instructions. The average size of each cDNA library was determined with a 2100 Bioanalyzer (Agilent Technologies, Santa Clara, CA, USA).

### Sequencing

cDNA libraries prepared as above were treated with an Ion PGM Sequencing 200 Kit (Life Technologies) and supplied to an Ion 318 Chip (Life Technologies) according to the manufacturer’s instructions. Sequencing was performed by using an Ion PGM next generation sequencer (Life Technologies). Sequencing data was subjected to the Maser analysis platform provided by National Institute of Genetics in Japan.

### Statistical analysis

Data were analyzed with one-way or two-way analysis of variance (ANOVA) and differences shown in ANOVA were analyzed with the Tukey’s method. The statistical analysis was also carried out by Student’s *t*-tests.

## Results

### Free amino acid content

The content of total free amino acids extracted from the abdominal muscle in the starting samples of kuruma shrimp (0 h) after rearing at 3.4 % salinity for three days was 260.7 ± 30.6 μmol g^-1^ tissue (Fig. 2). The most abundant free amino acid was glycine, followed by arginine, glutamine and alanine in most cases. ANOVA analysis revealed that the content of total free amino acids at 3.4 % salinity did not change significantly during rearing for another three days (72 h) (*P* > 0.05). The average content in the samples reared for 24 h at 1.7% salinity was significantly lower than that at 4.0 % salinity as well as that at 3.4 % salinity (Student’s *t*-test, *P* < 0.05). After 72 h, the content of total free amino acids at 4.0 % salinity was significantly higher than those at 1.7 % salinity (*P* < 0.01) and at 3.4 % salinity (*P* < 0.05). The content at 3.4 % salinity was also significantly higher than that at 1.7 % salinity (*P* < 0.05). The content of total free amino acids was also higher at 3.4 % than at 1.7 % salinity after rearing for 6 h (*P* < 0.05).

**Fig. 2.**
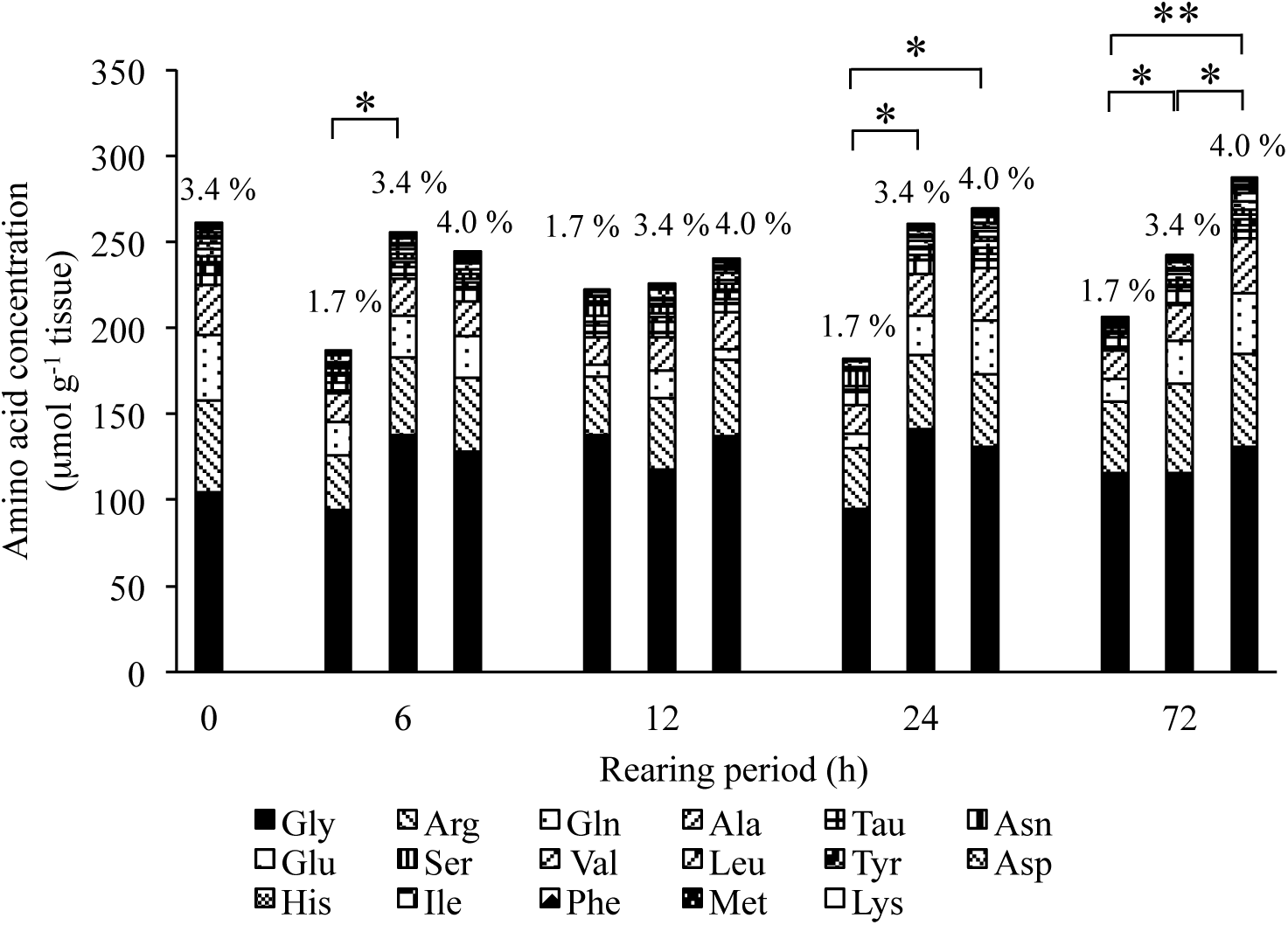
The concentrations of free amino acids in the second abdominal segments of kuruma shrimp reared at 1.7 %, 3.4 % and 4.0 % salinity for various rearing periods up to 72 h. Significance by Student’s *t*-test at \**P* < 0.05, \*\**P* < 0.01.

The changes in the major free amino acids during rearing were compared individually. No statistical differences were observed for glycine at any salinity (Fig. 3), although the content of glycine was the highest among all free amino acids (Fig. 2). The content of arginine having the second largest content showed significant difference only between the samples at 1.7 % and 4.0 % salinity after 72 h (Fig. 4).

**Fig. 3.**
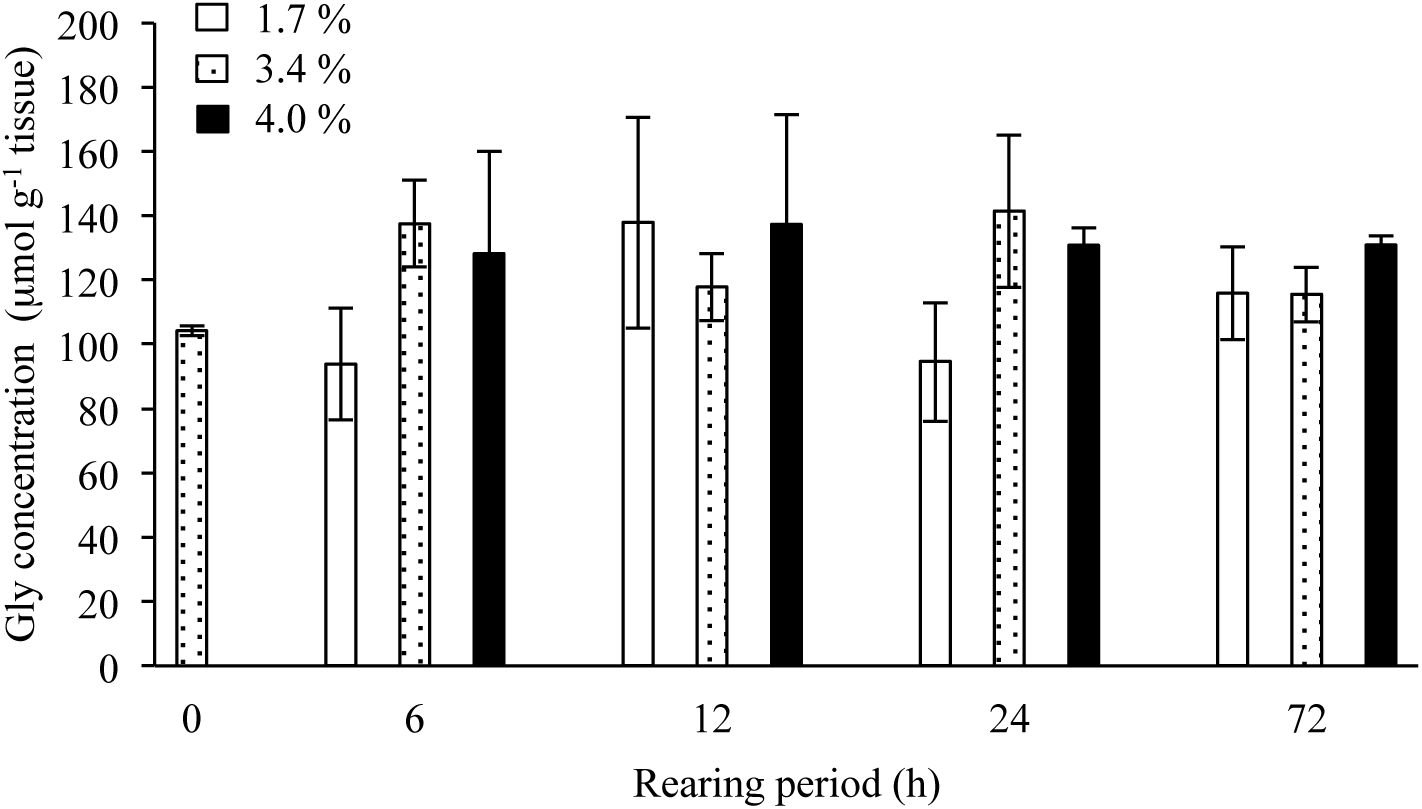
The concentration of glycine in the second abdominal segments of kuruma shrimp reared at different salinity for various periods up to 72 h. Open, dotted and solid bars indicate the concentrations of glycine in the samples reared at 1.7 %, 3.4 % and 4.0 % salinity, respectively.

**Fig. 4.**
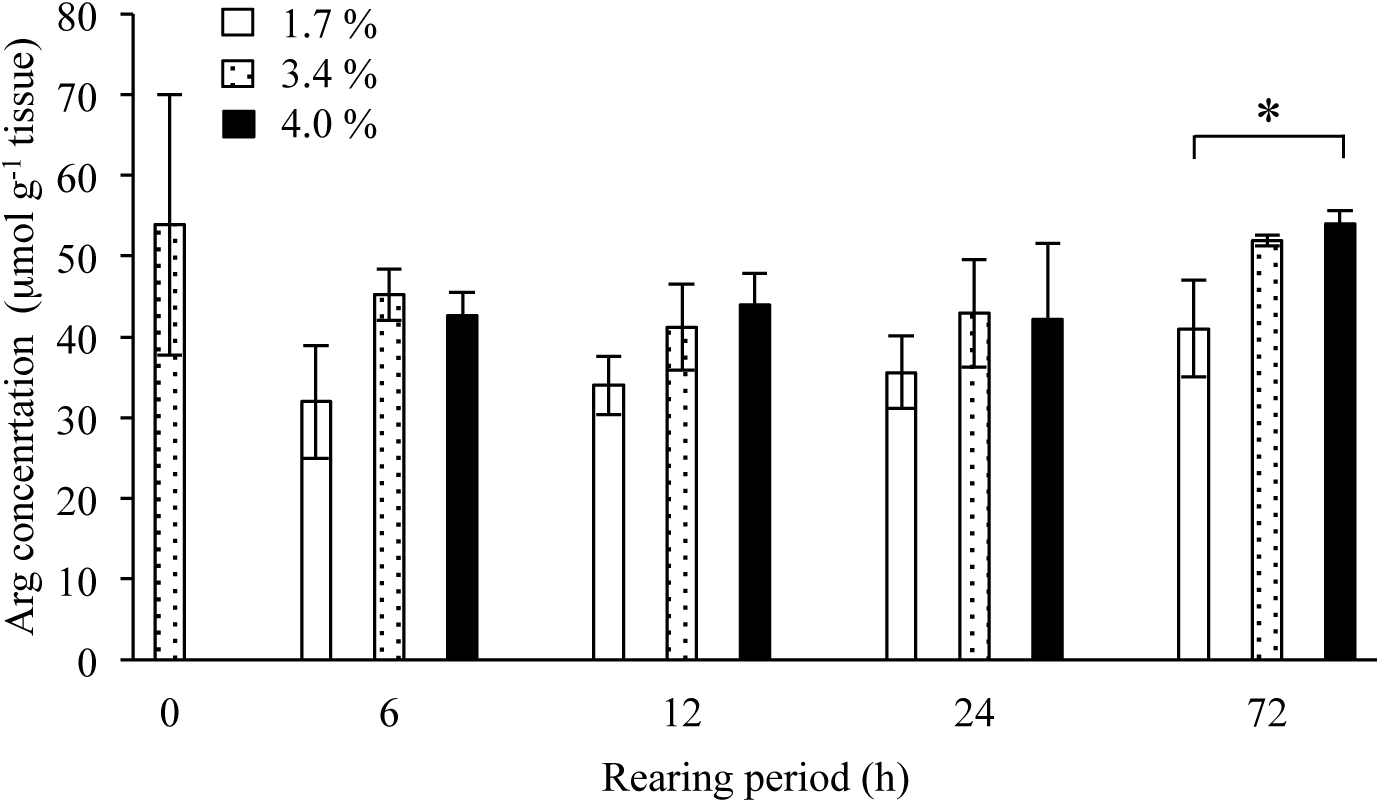
The concentration of arginine in the second abdominal segments of kuruma shrimp reared at different salinity for various periods up to 72 h. Open, dotted and solid bars indicate the concentrations of arginine in the samples reared at 1.7 %, 3.4 % and 4.0 % salinity, respectively. Significance by Student’s *t*-test at \**P* < 0.05.

Marked changes were observed in the content of glutamine as shown in Figure 5. The statistical analysis did not show any significant differences for the samples at 3.4 % salinity for any rearing periods, although the content for 0 h was apparently higher than those for 6 – 72 h. Student’s *t*-test revealed that the content at 4.0 % salinity was significantly higher than that at 1.7 % salinity after rearing for 24 and 72 h (*P* < 0.01). The content at 3.4 % salinity was also significantly higher than that at 1.7 % salinity after 72 h. ANOVA analysis demonstrated that the content at 1.7 % salinity after 6 h was significantly higher than that after 12 h, whereas the content at 4.0 % salinity after 12 h was significantly lower than those after 6, 24 and 72 h at the same salinity.

**Fig. 5.**
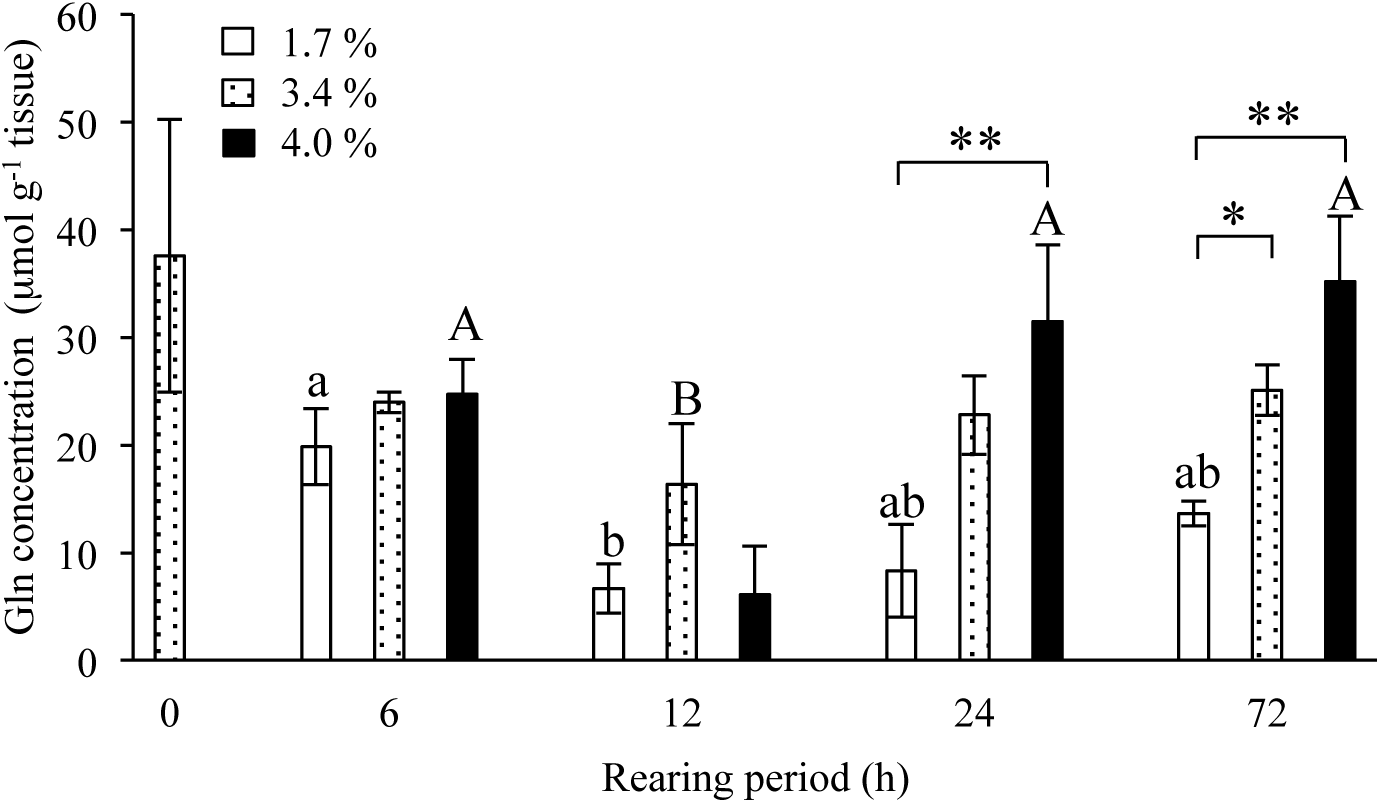
The concentration of glutamine in the second abdominal segments of kuruma shrimp reared at different salinity for various periods up to 72 h. Open, dotted and solid bars indicate the concentrations of glutamine in the samples reared at 1.7 %, 3.4 % and 4.0 % salinity, respectively. Significance by Student’s *t*-test at \**P* < 0.05, \*\**P* < 0.01. Different letters indicate significant differences by ANOVA.

The content of alanine showed the changes similar to those of glutamine as shown in Figure 6. The highly significant differences (*P* < 0.01) were observed in the content between the samples at 1.7 % and 4.0 % salinity after 24 and 72 h as well as between those at 3.4 % and 4.0 % salinity after 72 h. The difference between the samples at 1.7 % and 3.4 % salinity was also significant (*P* < 0.05). ANOVA analysis demonstrated that the contents after 24 and 72 h were significantly higher than those after 6 and 12 h for the samples at 4.0 % salinity. Taken together, it was found that the contents of glutamine and alanine changed clearly after rearing kuruma shrimp at 1.7 % and 4.0 % salinity.

**Fig. 6.**
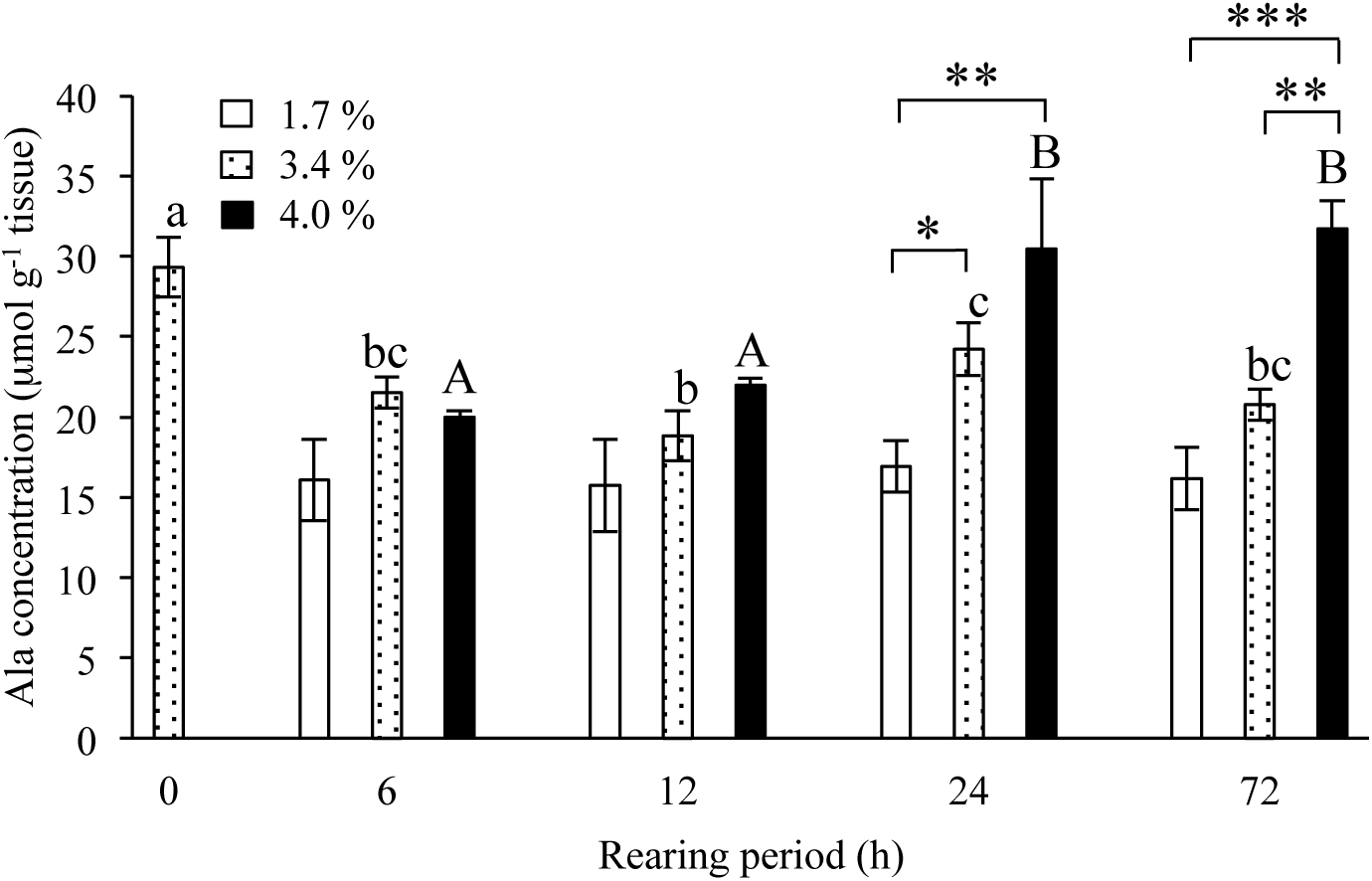
The concentration of alanine in the second abdominal segments of kuruma shrimp reared at different salinity for various periods up to 72 h. Open, dotted and solid bars indicate the concentrations of alanine in the samples reared at 1.7 %, 3.4 % and 4.0 % salinity, respectively. Significance by Student’s *t*-test at \**P* < 0.05, \*\**P* < 0.01, \*\*\**P* < 0.001. Different letters indicate significant differences by ANOVA.

### Water content

Figure 7 shows the changes in the water content of kuruma shrimp reared at different salinity. The water content was 74.9 % initially. ANOVA analysis demonstrated that the water content was not significantly changed when the samples were reared during 72 h at 3.4 % and 4.0 % salinity. On the other hand, the water content for samples reared for 24 h at 1.7 % salinity was significantly higher than those after 6 and 72 h. The differences in the water content for the samples after rearing for the same period at different salinity were significant between those at 1.7 % and 4.0 % salinity after 6 h (*P* < 0.01), 12 h (*P* < 0.05), 24 h (*P* < 0.01) and 72 h (*P* < 0.01). The significant differences were also observed between the samples at 1.7 % and 3.4 % salinity after 12 h, 24 h and 72 h (*P* < 0.05). However, no significant difference was observed between the samples at 3.4 % and 4.0 % when the contents after the same period were compared.

**Fig. 7.**
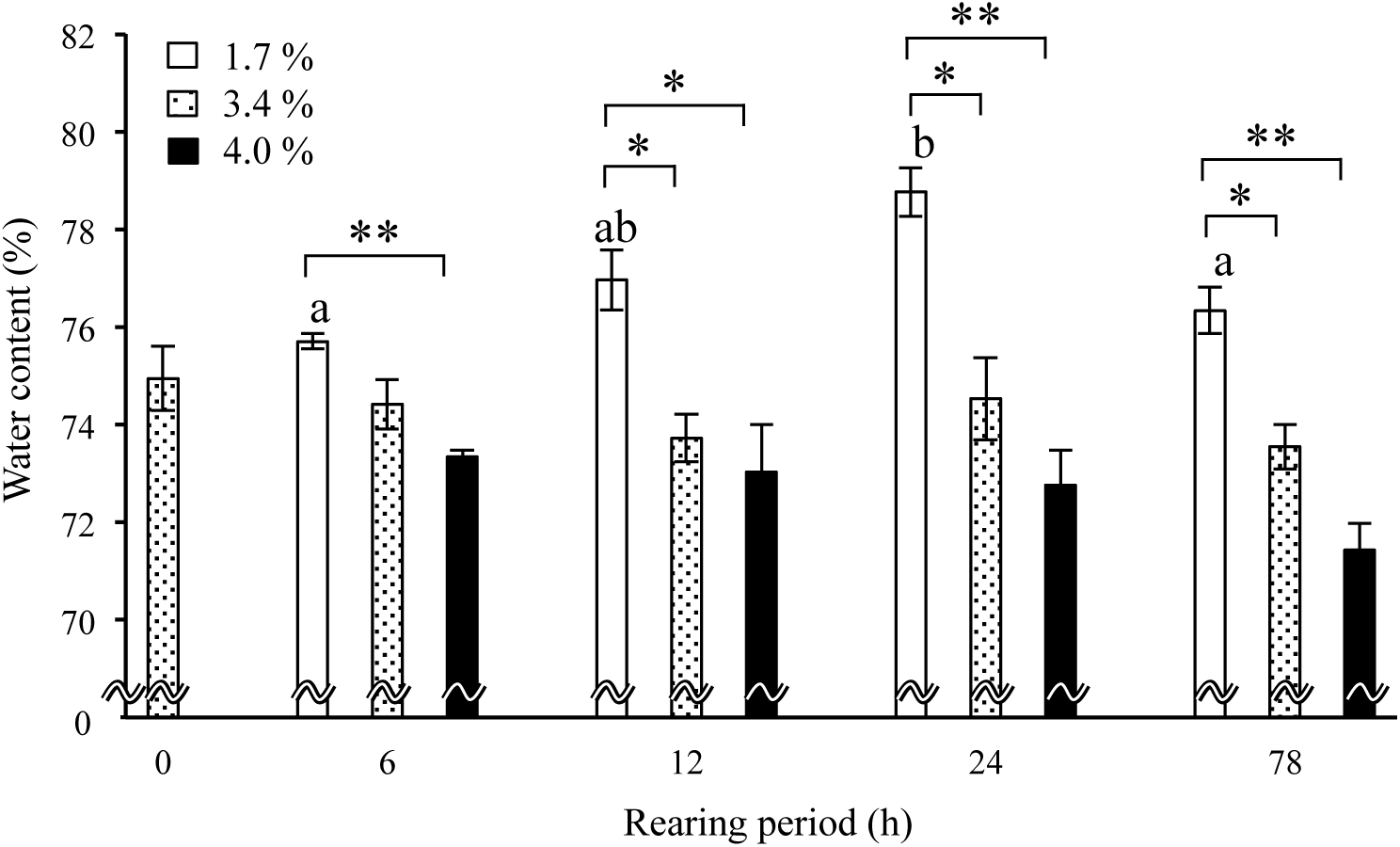
The water contents in the fourth and fifth abdominal segments of kuruma shrimp reared at different salinity for various periods up to 72 h. Dotted, dashed and solid bars indicate the water content in the samples reared at 1.7 %, 3.4 % and 4.0 % salinity, respectively. Significance by Student’s *t*-test at \**P* < 0.05, \*\**P* < 0.01. Different letters indicate significant differences by ANOVA.

Figure S1 shows the relationship between the water content and total free amino acid contents. The contents of total free amino acids were inversely proportional to the water content with a significant relative coefficient value of r = -0.54076.

### RNA-seq analysis

To minimize any possible individual variations, we mixed mRNAs prepared each from three specimens with the same amount (30 μg) for all sampling points as described in Materials and methods. Table S1 shows the average sizes of the constructed cDNA libraries for the samples reared for 24 h at 1.7, 3.4 and 4.0 % salinity, together with corresponding Ion PGM sequencing data. RNA-sequencing (RNA-seq) data for the samples reared at 1.7 % and 4.0 % salinity were subjected to the MA plot analysis (Wang *et al.*, 2010), together with those for the samples reared at 3.4 % salinity as a reference. The genes whose expression levels at 1.7 % salinity were increased and decreased more than 2 folds than those at 3.4 % salinity were 8,696 and 5,367, respectively (Table S2, Fig. S2). On the other hand, the corresponding numbers at 4.0 % salinity compared with those at 3.4 % salinity were 3,407 and 3,683, respectively (Table S2, Fig. S3).

Tables S3 - S6 show the genes whose expression levels were increased or decreased more than 10 folds during rearing for 24 h at 1.7 % or 4.0 % salinity compared with those at 3.4 % salinity as a reference, together with gene names, fragments per kilobase of transcript per million fragments (FPKM) values and fold change (FC). The genes whose expression levels were increased at 1.7 % salinity were 2,065, among which 124 genes were identified (Table S3). On the other hand, the genes whose expression levels were decreased at 1.7 % salinity were 2,618, among which 113 genes were identified (Table S4). Meanwhile, the numbers of the genes whose expression levels were increased and decreased at 4.0 % salinity were 444 and 302, respectively, among which 16 and 31 genes were identified, respectively (Tables S5 and S6).

### Gene expression profiles in glutamine- and alanine-related metabolic pathways

Glutamine-related metabolic pathways are shown in Fig. 8, where the changes in the expression levels of the genes encoding glutamate synthase, glutamate-ammonia ligase, glutaminase, glutamine-fructose-6-phosphate transaminase and amidophosphoribosyltransferase determined by RNA-seq analysis for the samples reared for 24 h at different salinity are depicted, together with the changes in the content of glutamine and glutamate after the same rearing period (Figs. 2 and 5). The expression levels of these genes except that encoding glutamate-ammonia ligase were decreased in association with the increase of salinity. On the other hand, the expression levels of the glutamate-ammonia ligase gene were increased following the increase of salinity.

**Fig. 8.**
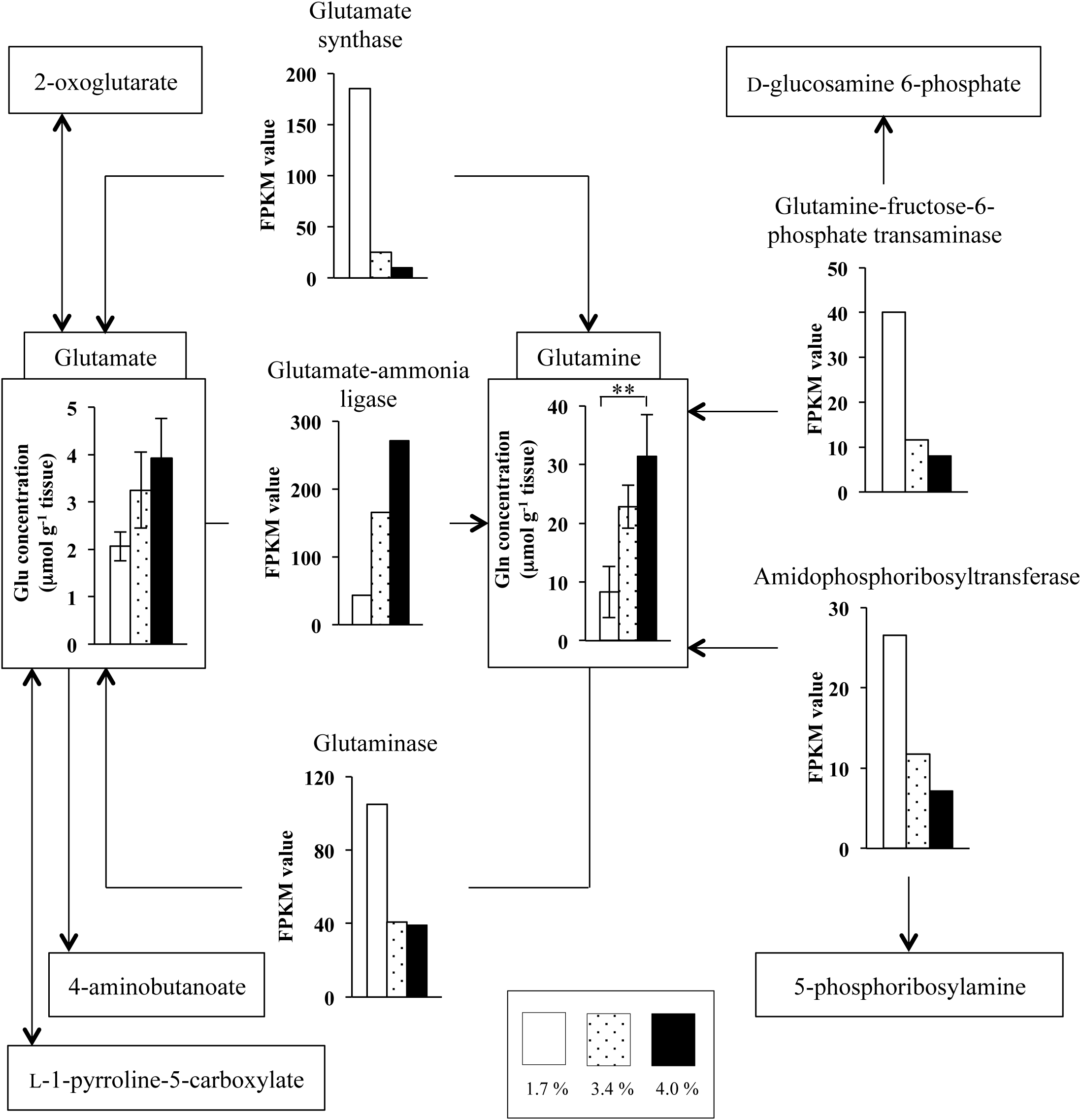
Genes and metabolites related to glutamine metabolism. Open, dotted and solid bars in squared panels indicate the contents of glutamine and its related metabolites in the samples reared for 24 h at 1.7 %, 3.4 % and 4.0 % salinity, respectively, whereas those not squared indicate the FPKM values of glutamine-related metabolic genes at corresponding salinity, respectively.

Alanine-related metabolic pathways are shown in Fig. 9, where the changes in the expression levels of the genes encoding alanine transaminase and alanine-glyoxylate transaminase determined by RNA-seq analysis for the samples after 24 h at different salinity as those for glutamine are depicted, together with the changes in the content of alanine after the rearing period of 24 h at different salinity (Figs 2 and 6). The expression levels of the alanine dehydrogenase gene were decreased following the increase of salinity, whereas those of the alanine-glyoxylate transaminase gene were higher for the samples at 3.4 % salinity than those at 1.7 % and 4.0 % salinity. Unfortunately, the gene encoding alanine dehydrogenase could not be detected in the present RNA-seq analysis.

**Fig. 9.**
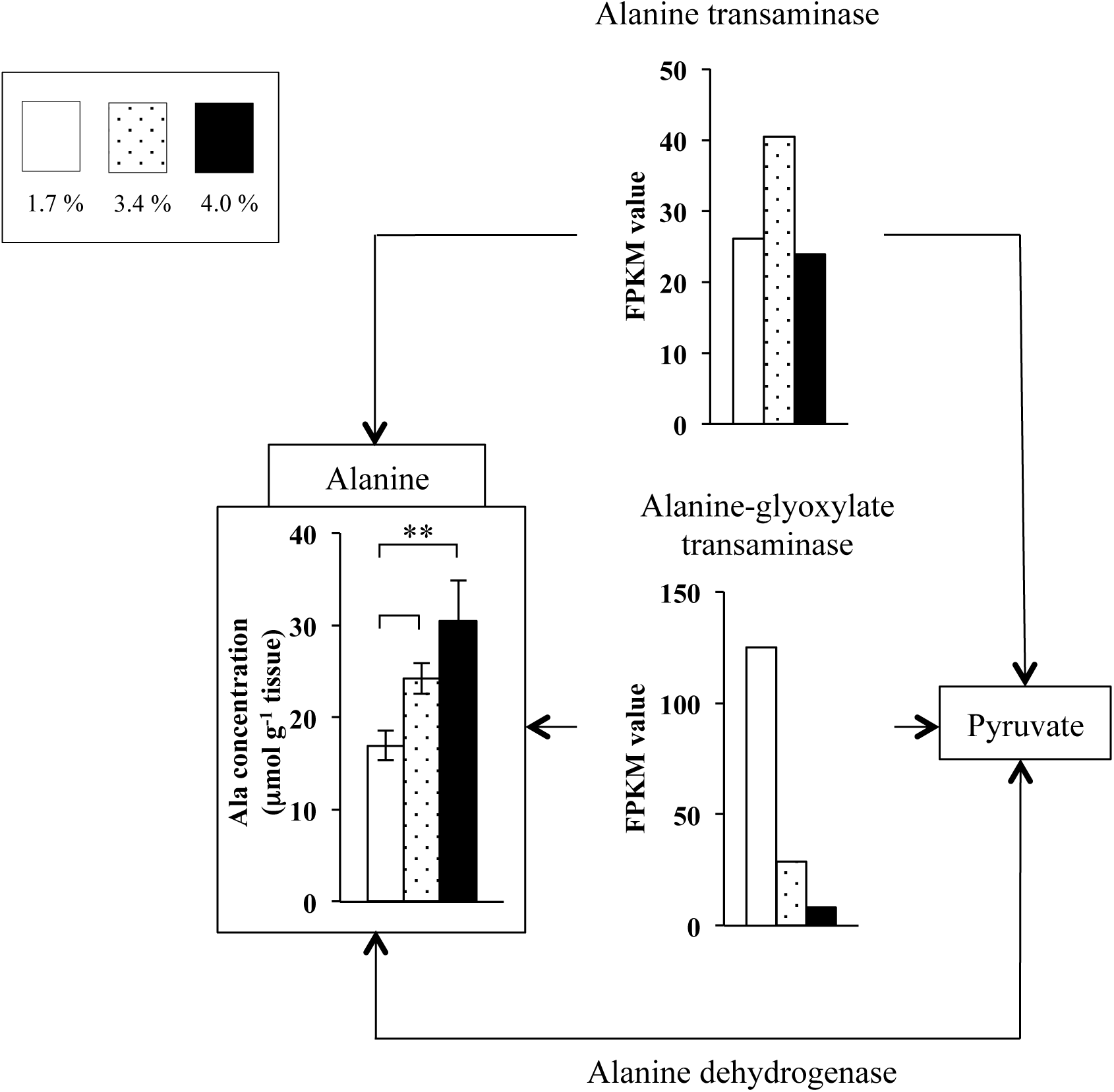
Genes and metabolites related to alanine metabolism. Open, dotted and solid bars in squared panel indicates the content of alanine in the samples reared for 24 h at 1.7 %, 3.4 % and 4.0 % salinity, respectively, whereas those not squared indicate the FPKM values of alanine-related metabolic genes at corresponding salinity, respectively.

### Expression profiles of genes other than those related to amino acid metabolism

RNA-seq analysis revealed that many genes other than those related to amino acid metabolism altered their expression levels, when the samples were reared for 24 h at different salinity (Tables S3 – S6, Figs. S2 and S3). The expression levels of the genes encoding myosin heavy chain (MYH) type 1 (MYH1) and MYH2 were increased at 1.7 % salinity (Table S3). The expression levels of MYH3 were decreased at 1.7 % salinity (Table S4) and increased at 4.0 % salinity (Table S5).

The expression levels of the genes encoding sarco/endoplasmic reticulum Ca^2+^-ATPase (SERCA) and ATP synthase subunit 9 mitochondrial precursor were decreased at 1.7 % salinity. On the other hand, the expression levels of the gene encoding Na^+^/K^+^-ATPase α-subunit were increased at 1.7 % salinity (Table S3).

## Discussion

In order to understand the mechanisms involved in the adaptation of kuruma shrimp to different environmental salinity, we determined the contents of free amino acids in the abdominal muscle from the samples reared at different salinity for various time periods until 72 h. As shown in Fig. 2, the contents of total free amino acids were changed following the alternation of rearing salinity. It has been reported that the concentrations of total free amino acids in kuruma shrimp were increased following the increase of salinity during 4 days (Abe *et al.*, 2005). We obtained similar results in the present study. In addition, the present study demonstrated that it took 24 h to change the contents of free amino acids following the alternation of salinity.

Figure 2 also shows that glycine, arginine, glutamine and alanine were the major free amino acids as reported previously (Abe *et al.*, 2005). Figures 3 - 6 show the changes in the contents of these major amino acids, respectively. The content of glycine did not change significantly (Fig 3), although previous investigations observed the increase of the glycine content in the muscles of kuruma shrimp and crayfish (Okuma and Abe, 1994; Abe *et al.*, 2005). The content of arginine neither changed significantly following the increase of salinity (Fig. 4), whereas those of glutamine and alanine were increased at high salinity and decreased at low salinity after rearing for 24 h (Figs. 5 and 6). Therefore, these two amino acids were found in the present study to possibly play as osmolytes, taking at least 24 h to adjust the cellular osmotic pressure to the environmental salinity. We reported previously the changes in the accumulation of metabolites in brackish water clam *Corbicula japonica* exposed to different salinity, where the content of L-alanine was also increased in association with the increase of environmental salinity, although that of L-glutamine was not changed significantly (Koyama *et al*., 2015; Okamoto *et al*., 2012). Therefore, it is considered that alanine is an osmolyte common to invertebrates having open vascular systems.

The water content of the samples reared at 1.7 % salinity was increased significantly during the rearing period up to 24 h and returned to the original level subsequently (Fig. 7). On the other hand, the water content in the samples reared at 4.0 % salinity tended to be decreased gradually during rearing for 72 h, although this tendency was not significantly by ANOVA.

The increase of water content was almost proportional to the decrease in the content of total free amino acids (r = -0.54076, Fig. S1). Although the contents of free amino acids were increased following the decrease of the water content, the difference in the water content between the samples reared for 72 h at 1.7 % and 4.0 % salinity (5 %) was much less than that in the content of total free amino acids between the same samples (30 %). Thus, the increase in the content of total free amino acids following the increase of salinity is not likely the simple effect of the decrease in the water content, but seems attributable to the adaptation of kuruma shrimp to high salinity by increasing the concentration of free amino acids as reported previously (Okuma and Abe, 1994; Abe *et al.*, 2005).

Figure 8 shows the metabolic pathways of glutamine and its related compounds. The content of glutamine was increased following the increase of salinity after rearing for 24 h (Fig. 5). The FPKM value of the gene encoding glutamate-ammonia ligase was also increased following the increase of salinity, and those of other four genes found in the present study were rather decreased at high salinity (Fig. 8). Therefore, it is considered that glutamine was synthesized from glutamate by glutamate-ammonia ligase in response to high salinity, thus increasing the concentration of glutamine at this salinity. Other four enzymes including glutamate synthase, glutaminase, glutamine-fructose-6-phosphate transaminase and amidophosphoribosyltransferase all enhanced their expression at 1.7 % salinity. These results suggest that glutamine were catabolized to glutamate, D-glucosamine 6-phosphate or 5-phosphoribosylamine, thus decreasing the concentration of glutamine at this low salinity.

Figure 9 shows the metabolic pathways of alanine and its related compounds. As in the case of glutamine, the content of alanine was increased following the increase of environmental salinity after 24 h (Fig. 6). The FPKM value of the alanine-glyoxylate transaminase gene was increased at low salinity, whereas that of the alanine transaminase was not changed markedly at different salinity. Thus it seems that alanine was catabolized to pyruvate by alanine-glyoxylate transaminase gene at low salinity, decreasing the concentration of alanine at this low salinity. However, we could not detect the transcripts encoded by the alanine dehydrogenase gene. Thus, the mechanisms involved in the changes of the alanine content at different salinity have remained unclear. In this regard, it has been reported that pyruvate was not detected in the gill and foot muscle of brackish water clam, although the content of alanine was increased at high salinity (Koyama *et al.*, 2015). Pyruvate once accumulated at low salinity might have been quickly catabolized into another substance.

Although we did not carry out real-time PCR to confirm the data obtained from global gene expression analysis by RNA-seq in the present study, we previously demonstrated on similar works for fish that the RNA-seq data were satisfactorily verified by real-time PCR experiments (Ikeda *et al*., 2017; Tan *et al*., 2012)

The expression levels of the MYH1 and MYH2 genes were increased at 1.7 % salinity (Table S3). MYH is the major muscle protein and two types of the MYH genes, *MHC1* and *MHC2*, which correspond to the genes encoding MYH1 and MYH2, respectively, have been reported to be expressed in the abdominal muscle of kuruma, black tiger and Pacific white shrimps (Koyama *et al.*, 2012a, b). These are expressed only in flexor muscle and both in flexor and extensor muscles having anaerobic metabolism, respectively. In the present study, the expression levels of MYH1 at 3.4 % salinity were about 1.5 folds more than that of MYH2, and the expression levels of MYH2 at 1.7 % salinity were about 3 folds more than that of MYH1. The expression levels of MYH3 were decreased at 1.7 % salinity (Table S4) and increased at 4.0 % salinity (Table S5). The MYH3 gene has been reported as *MHC3* to be expressed in the pleopod muscle having aerobic metabolism of kuruma shrimp and black tiger shrimp (Koyama *et al.*, 2013).

SERCA plays an important role to regulate the calcium concentration in cytoplasm (Clapham, 1995). It has been reported that the expression levels of SERCA in the muscle of Pacific white shrimp reared at high salinity were higher than those at low salinity (Wang *et al*., 2013). Although, the expression levels of the SERCA gene were decreased at 1.7 % salinity, we did not observe the increase at 4.0 % salinity in this study (Table S4). The expression levels of the ATP synthase subunit 9 mitochondrial precursor gene were also deceased at 1.7 % salinity (Table S4). Such decreased expression levels of this gene have been reported in the gill from Pacific white shrimp reared at low salinity (Gonçalves-Soares *et al.*, 2012).

The expression levels of the Na^+^/K^+^-ATPase α-subunit gene were increased at 1.7 % salinity (Table S3). Such enhanced expression of the Na^+^/K^+^-ATPase α-subunit gene has been reported in the gill from black tiger shrimp reared at low salinity (Shekhar *et al.*, 2013), suggesting that ion transporters such as Na^+^/K^+^-ATPase are considered to play an important role to adjust the cellular osmotic pressure to the environmental salinity.

In conclusion, we examined the contents of free amino acids and gene expression profiles of kuruma shrimp reared at different salinity. A number of genes changed their expression levels to adapt the environmental salinity. In addition, the contents of free amino acids were considered to be regulated by various genes related to amino acid metabolism. For instance, the content of glutamine was increased at high salinity in association with the increase of the expression level of glutamate-ammonia ligase. The content of alanine was increased at high salinity in association with the decrease of the expression level of alanine-glyoxylate transaminase. In the future study, we need further investigation including the participation of D-amino acids and their related enzymes in the adaptation of shrimps to environmental salinity change.

## Author contributions

H. K. and S. W. conceived the study; H. K., N. M., M. H., E. T., D. I. and T. Y. designed experiments and collected the data; H. K., N. M. and S. W. analyzed data; H. K. and S. W. wrote the paper; H. K., N. M., K. Y., M. J. and S. P. interpreted the data; H. K., N. M. and S. W. contributed substantially developing the manuscript and take full responsibility for the content of the paper.

## Data availability

RNA-seq data are available in the DDBJ database under the accession number of DRA 006082.

## Supplementary information

Supplementary information available online at http://jeb.biologists.org/

## List of symbols and abbreviations

ANOVA: analysis of variance
DDBJ: DNA Data Bank of Japan
FPKM: fragments per kilobase of transcript per million fragments
FC: fold change
gDNA: genomic DNA
MHC: myosin heavy chain
MYH: myosin heavy chain
RNA-seq: RNA-sequencing
SERCA: sarco/endoplasmic reticulum Ca^2+^-ATPase

## Figure legends

Fig. S1. The relationship between water content and total free amino acid content. The relationship significant with coefficient of correlation (r) of -0.54076. Significance by test for non-correlation at \*\**P* < 0.01.

Fig. S2. MA plot constructed from RNA-seq data from kuruma shrimp reared for 24 h at different salinity. The plot shows the comparison of the gene expression levels of the samples at 1.7 % salinity with those at 3.4 %. The x-axis represents the average log value of the gene expression levels (A) and y-axis represents the logarithmic value of the fold changes (FC) in the gene expression levels (M). Diamonds correspond to individual genes. Black diamonds represent the genes whose expression levels at 1.7 % salinity were increased or decreased less than 2 folds than those at 3.4 %. Red and blue diamonds represent the genes whose expression levels at 1.7 % salinity were increased and decreased more than 2 folds than those at 3.4 %, respectively.

Fig. S3. MA plot constructed from RNA-seq data from kuruma shrimp reared for 24 h at different salinity. This plot shows the comparison of the gene expression levels of the samples at 4.0 % salinity with those at 3.4 %. Refer to the legend of Fig. S2 for symbols and x- and y-axes.

